# Woodsmoke and Diesel Exhaust: Distinct Transcriptomic Profiles in the Human Airway Epithelium

**DOI:** 10.1101/2025.02.26.640436

**Authors:** Ryan D. Huff, Christopher F. Rider, Theodora Lo, Kristen I. Hardy, Nataly El-Bittar, Min Hyung Ryu, Chris Carlsten, Emilia L. Lim

## Abstract

Climate change is increasing the frequency and severity of wildfires globally, causing significant woodsmoke (WS) emissions. Vehicles emit sizable amounts of toxic traffic-related air pollution (TRAP), for which diesel exhaust (DE) is a model. Both WS and DE contain particulate matter <2.5 microns (PM_2.5_) which deeply penetrates the lungs, causing respiratory epithelial inflammation that drives health effects. Regulations focus on PM_2.5_ concentration, despite emerging research that highlights how composition mediates health effects. As WS and DE are compositionally distinct, we conducted the first head-to-head comparison of effects on the transcriptomes of air-liquid interface cultured primary human bronchial epithelial cells (HBEC). Differentiated donor-matched HBEC transwells were exposed for 2-hours to filtered air (FA; control), or WS (furnace tube burning pine) or DE (Hatz 1B30E generator) both diluted to 300 µg/m^3^ of PM_2.5_. WS had higher ultrafine PM, whereas DE exposure contained significantly higher NO_2_, CO, and O_3_. RNA sequencing showed that WS exposure resulted in 119 (↑41, ↓78) differentially expressed genes, while DE modulated 399 (↑255, ↓144) compared to FA exposure. WS was associated with small ribosomal subunit and cytochrome complex related genes, while DE exposure was associated with HIF-1 signaling, respiratory chain complex and interferon alpha/beta signaling/ISG15-protein conjugation, suggesting how TRAP exposure may enhance infection risk. We also analyzed exposure effects on protein immune-mediators. We demonstrate that two major air pollution sources modulate different genes and pathways in HBECs, with minimal overlap. This informs the debate regarding the regulatory focus on concentration and assumptions that similar concentrations of air pollution have indistinct effects.

**Highlights:**

- Lung health effects of diesel exhaust (DE) and wood smoke (WS) are underexplored
- RNAseq of DE and WS-exposed primary lung epithelial cells revealed larger DE effects
- DE showed greater repression of host antiviral response-associated genes than WS
- A regulatory focus on PM concentration may miss composition-specific lung effects

## 1. Introduction

Climate change is increasing the frequency, severity, and size of wildfires globally. Large-scale landscape fires emit millions of tons of smoke, composed of toxic gases and particulate matter (PM) that can cause widespread episodes of reduced air quality (Byrne et al., 2024). Vehicles also emit a sizable portion of gases and PM, together known as traffic-related air pollution (TRAP). Most governmental regulations have assumed that only PM concentration is relevant, despite emerging research demonstrating health effects are considerably affected by composition (Weichenthal et al., 2024). Indeed, PM less than 2.5 microns in diameter (PM_2.5_) generated from wildfires may be more harmful than ambient urban air pollution, as it is associated with increased hospitalizations in Southern California (Aguilera et al., 2021).

The combustion of fossil or biomass fuels releases primary aerosols into the atmosphere that include elemental and organic carbon, and metals such as aluminum (Al), iron (Fe), zinc (Zn), copper (Cu), nickel (Ni), and manganese (Mn). The composition of these emissions is impacted by the type, source and quality of fuels, engine characteristics in the case of TRAP, temperature, and humidity (Al-Kindi et al., 2020; Heydarzadeh et al., 2022). Furthermore, secondary organic aerosols are produced by the oxidation of emitted precursors, and consist of sulfur, nitrogen, and organic carbon species (including volatile organic compounds (VOCs) and polycyclic aromatic hydrocarbons (PAHs)) at varying levels (Al-Kindi et al., 2020; Kelly and Fussell, 2012).

TRAP, which can be modelled by diesel exhaust (DE), is typically characterized by increased concentrations of carbon monoxide (CO), sulfur oxides (SO_x_), nitrogen oxides (NO_x_), and ozone (O_3_), as well heavy transition metals such as Ni, Fe, Zn, and Mn (Yin et al., 2020). Whereas increased concentrations of CO and nitric oxide (NO) gases, and increased fractional concentrations of magnesium (Mg), phosphorus (P), and potassium (K) have been measured during wildfire episodes. Wildfire smoke also contains increased levels of volatile organic compounds (VOCs), yet often has low NO_x_ levels (Verma et al., 2009). Furthermore, it is known that PM particles released by wildfire emissions are smaller and predominately fall within the ultrafine or PM_1_ particle size range (<2.5 microns in diameter) (Sparks and Wagner, 2021). However, a significant portion of the PM in both wildfire smoke and DE falls in the PM_2.5_ range, a size of particle known to deeply penetrate the lungs and contribute to a wide range of acute and chronic health effects (Thangavel et al., 2022).

Fire smoke is a broad term for emissions released from the combustion of biomass and includes woodsmoke (WS) and wildfire smoke. It is important to note that wildfire smoke contains a much more diverse set of chemicals than woodsmoke, as the smoke can be generated from the combustion of a range of both natural and man-made fuels. However, woodsmoke contains many of the same particulates and VOCs as wildfire plumes and the health impacts have been studied in detail (O’Dell et al., 2020; Singh et al., 2023). The direct effect of woodsmoke on the human lungs has previously been investigated in over ten controlled human exposure studies to date (Schwartz et al., 2020). Notably, these studies indicate acute exposure to wood smoke (< 4 hours) may be associated with lung and systemic inflammation. Similar results have been observed in acute exposure studies to DE (Huff et al., 2019; Long et al., 2022; Orach et al., 2021). Despite this research, little is known about how the compositional similarities and differences between WS and DE impact human health, as no direct comparison has been performed (Long et al., 2024).

The pseudostratified lung epithelium, along with resident and infiltrating immune cells, represents the primary barrier to exogenous insults such as WS and DE in the lungs. As the first line of defence, the epithelium detects xenobiotic material and orchestrates subsequent immune responses (Hewitt and Lloyd, 2021). *In vitro* models of the lung epithelium have identified similar molecular effects of WS and DE exposures on the epithelium, which include increased production of mucin proteins, barrier dysfunction, pro-inflammatory signalling, oxidative stress, and macromolecular damage and toxicity (Huff et al., 2019; Memon et al., 2020; Roscioli et al., 2018; Upadhyay et al., 2024; Zeglinski et al., 2019). Despite these similarities, little is known about the differential cellular effects of WS and DE, as direct comparisons of live aerosol exposures to WS or DE within the same model system have not been performed (Hewitt and Lloyd, 2021; Memon et al., 2020; Roscioli et al., 2018; Wang et al., 2021; Zeglinski et al., 2019).

As WS and DE are compositionally distinct, in the present study we conducted the first head-to-head comparison of the effects of live aerosol exposures on the transcriptomes of air-liquid interface (ALI) cultured primary human bronchial epithelial cells (HBEC). RNA-sequencing (RNA-seq) captures the cellular response to each of these exposures. This approach allows in-depth evaluation of the impact of WS and DE exposures on the human airway epithelium. Using this approach, we tested the hypothesis that DE and WS induce a distinctive cellular response in the airway epithelium. Furthermore, we assessed several characteristics of freshly generated WS and DE in real-time when the PM_2.5_ mass concentration was kept consistent at 300 µg/m^3^. We also assessed the corresponding effect of these exposures on the transcriptome and immune mediator profiles of HBECs (Figure 1). Our findings add new evidence supporting the notion that similar concentrations of combustion-related air pollution have distinct effects depending on the emission source. This may inform the debate regarding whether ambient pollution should be regulated based on composition, as well as concentration (Weichenthal et al., 2024).

**Figure 1.**
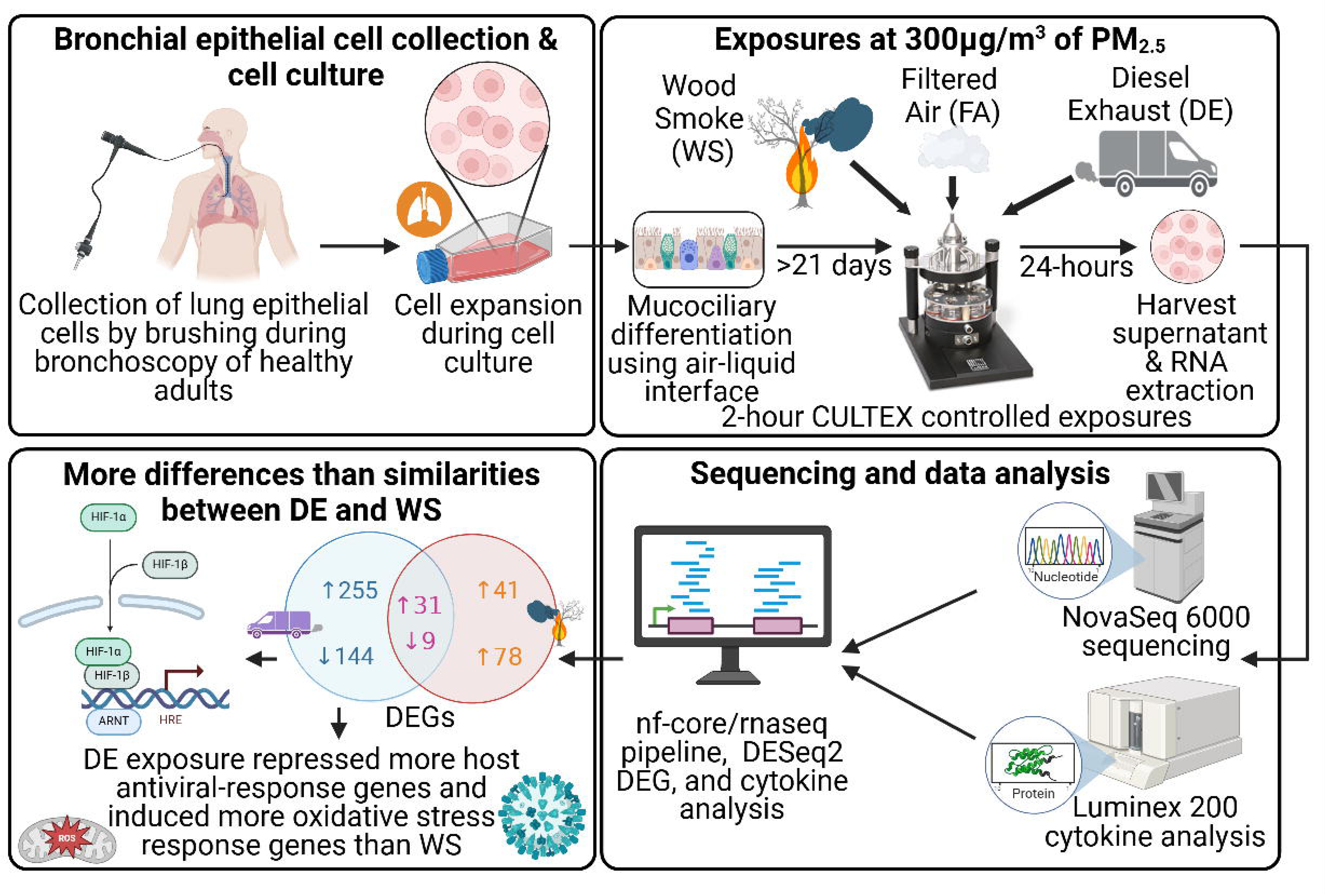
Study overview and key findings. Human bronchial epithelial cells were collected from a bronchoscopy study. Cells were then cultured and differentiated under airway-liquid interface conditions *in vitro*. Cells were subsequently exposed directly to fresh diesel exhaust (DE) or wood smoke (WS) at an equal 300µg/m^3^ particle mass concentration using a Cultex exposure system. Cells and supernatants were harvested 24 hours after exposure. RNA-sequencing and Luminex 200 cytokine analysis were performed to analyze the epithelial response to each pollutant. Created in BioRender.

## 2. Methods

### 2.1 Clinical study background

HBECs used in the study were collected from two clinical bronchoscopy studies: the “Air, Gene, and Environment Study” (AGnES) (Tiew et al., 2024) and “COPD originates in polluted air” (COPA; controlled human DE exposure study) (Ryu et al., 2022). Both studies received approval from the ethical review boards of the University of British Columbia and the Vancouver Coastal Health Research Institute, as H18-01836 and H14-00821, respectively. For the current project, we included HBECs from healthy control participants without a diagnosis of COPD. Participants gave written informed consent prior to undergoing research bronchoscopies. Bronchial brushings, collected from the endobronchial mucosa of a 4-5^th^ order airway in 3 male and 3 female healthy adults, were used in this study (Table E1).

### 2.2 Human bronchial epithelial cell culture

HBECs were expanded from the cells collected on endobronchial brushings in PneumaCult-EX Basal Medium (Stemcell Technologies, Vancouver, BC) and passaged once before cryopreservation. Cryopreserved cells were thawed and expanded in T-25 cell culture flasks denoted P2 (passage 2). Once expanded to ∼80% confluence, HBEC were seeded onto 6.5 mm transwell polyester membrane inserts with a 0.4 μm pore size (Corning) at a concentration of 3×10^4^ cells/insert (0.33 cm^2^ growth area). HBECs were then incubated in submerged cultures for 4 days in PneumaCult-EX Basal Medium, prior to air lifting using PneumaCult-ALI Maintenance Medium (Stemcell Technologies) as per the manufacturer’s instructions. Cells were then maintained under ALI culture conditions for at least 21 days prior to experiments. Trans epithelial electrical resistance (TEER) was checked to make sure air-liquid barrier was formed. TEER was measured using a Millicell ERS-2 Voltohmmeter (EMD Millipore, Canada). Mucus was washed from the apical surface every 2-3 days using Hanks’ Balanced Salt Solution (HBSS), including on the day prior to exposures.

### 2.3. Diesel exhaust and wood smoke exposures

DE was freshly generated using a Hatz 1B30E generator (United States Environmental Protection Agency (EPA) Tier 4/European Union Stage 5 compliant) (Birger et al., 2011; Orach et al., 2023, 2022). WS was also freshly produced using a US EPA-designed furnace tube (Kim et al., 2019, 2018; US EPA, 2019). Lodgepole pine (*Pinus contorta),* an international standard wood type sourced from the EPA, was burned using a computer-controlled ceramic heating element that can reach over 1000°C (Kim et al., 2018). Lodgepole pine is prevalent in Western Canadian forests (and, therefore, wildfires) and is a tree of choice for Canada’s largest national firewood distributors, and used in BC-produced bio pellets shipped around the world (Bull and Bennett, 2022; Gagné et al., 2017; Ministry of Forests, 2024).

Both DE and WS were diluted to 300μg/m^3^ PM_2.5_ using high-efficiency particulate air (HEPA)-filtered air, with the majority of smoke generated directed into piping, taking it out of the building without measurement. Subsequently, the diluted aerosol flowed into the exposure booth, which contains extensive particulate matter and gas monitoring equipment (see Table E2), and then a CULTEX *in-vitro* radial flow system (Aufderheide et al., 2013; Buckley et al., 2024). The Cultex was run according to manufacturer’s specifications, exposing cells at a rate of 5 mL per min per insert for two hours. Differentiated HBECs in participant-matched transwells were kept in the incubator (IC; control), or exposed for 2-hours to HEPA and charcoal-filtered air (FA; control), or DE (Hatz 1B30E generator) or WS (furnace tube burning lodgepole pine), with DE and WS exposure completed on the same day, at the Air Pollution Exposure Lab (Birger et al., 2011). Basolateral supernatants were collected at 24-hours for cell viability analysis by lactate dehydrogenase (LDH), and multiplex cytokine assays. Cells were harvested for RNA sequencing 24-hours after exposure. Total RNA was extracted using RNEasy plus kits (Qiagen), frozen and sent on dry ice for sequencing at the Michael Smith Genome Science Centre.

### 2.4. Cell protein assays

Supernatants were collected from the basolateral compartment of transwell ALI inserts 24-hours after the start of the exposure. Analyte concentrations (pg/mL) from undiluted samples were determined using a Human Cytokine 96-Plex Discovery Assay performed by Eve Technologies (Calgary, Alberta, Canada). Analytes found to be undetectable were eliminated from the analysis; otherwise for statistical comparisons values found to be below the lower limit of detection (LLOD) were replaced with 1⁄2 of the respective LLOD.

Lactate dehydrogenase (LDH) assays were performed using a CyQUANT LDH cytotoxicity assay kit (Thermo-Fisher Scientific, USA) as per the manufacturer’s instructions using basolateral supernatants. Absorbance was measured using a Spectramax i3x plate reader (Molecular Devices, USA). TEER was measured 24-hours post-exposure and multiplied by insert growth area (Ohms*cm^2^). LDH and TEER data were analyzed by one-way ANOVA with Tukey’s post-hoc test (n = 6). Both LDH (Figure E1A) and TEER (Figure E1B) were not significantly different between controls and exposures.

### 2.5. Sequencing and data analysis

A detailed protocol for the poly-A tail enrichment and cDNA synthesis is described in our online supplement. RNA sequencing was performed with an Illumina NovaSeq 6000 sequencer targeting 50M read-pairs per library. RNA-seq reads were quality checked and pre-processed using the nf-core/rnaseq pipeline (v3.14.0-gb89fac3), using the GrCH38 human reference genome and gene annotation as inputs. Patient-matched FA samples were merged as technical replicates, and Salmon quasi-mapping used for transcript quantification.

DESeq2 v1.42.1 was used in R v4.3.2 to perform differential gene expression analysis with the model “∼participant + basalCounts + condition”, which adjusted for the normal variation in the proportions of basal cells in each sample (Figure E2), and the contrast c(“condition”, {“DE”, “WS”}, “FA”). Pathway analysis was subsequently performed on the differentially expressed genes (DEGs) using clusterProfiler v4.10.11.

To adjust for the cell population composition (basalCounts) in our differential gene expression analysis, CIBERSORTx (https://cibersortx.stanford.edu), calibrated on Human Lung Cell Atlas (HLCA) (Sikkema et al., 2023) data, was used to impute the proportions of basal and other epithelial cell types from the count matrix values (CPM values) of each sample (Figure E2) (Newman et al., 2019). Our GitHub page (https://github.com/EmiliaLimLab/DE_WS_RNAseq) contains the custom signature matrix. The raw gene counts matrix was counts per million (CPM) transformed and any genes linked to mitochondria or missing HGNC gene symbol annotation were removed. Genes that were not expressed (CPM value = 0) in >80% of samples were also filtered. Therefore, 21,971 genes were used. The signature matrix was trained on the HLCA dataset. For this we removed any data from active smokers. Cells annotated as “epithelial cells” were used. We excluded all nasal epithelium cells from training. We downsampled the data to 1000 for each level 3 annotation (ann_level_3). CIBERSORTx was run on the Stanford server with default parameters for single cell RNA-seq data signature matrix generation. Quantile normalization was disabled. Kappa=999, q-value = 0.01, number of barcode genes set to 300-500, min expression = 1, replicates = 5, sampling set to 0.5, and filter for non-hematopoietic genes set to false. Subsequently, we performed deconvolution in fraction mode with the number of permutations set to 100 and absolute mode set as equal to true.

### 2.6. Network analysis

Protein-protein interaction network analysis was performed using the DEGs using STRINGdb (https://string-db.org) version 12.0 on Feb 11, 2025. Ensemble gene ids of DEGs were queried on the multiple proteins search. The interaction sources were “Experiments”, “Databases”, “Co-expression”, “Neighborhood”, “Gene Fusion”, and “Co-occurrence”. We included the full STRING network, where the edges indicate both functional and physical protein associations.

## 3. Results

### 3.1. Effects of diesel exhaust and wood smoke exposures on gene expression

Despite having overlapping PM_2.5_ concentrations, WS (306.5±107.2 µg/m^3^) had a higher proportion of ultrafine PM, whereas DE (322.3±82.9 µg/m^3^) had significantly higher levels of NO_2_, CO, O_3_, and total volatile organic compounds (Table E2). Even so, principal component analysis of our RNA-sequencing data showed that samples clustered by donor, rather than exposure (Figure 2A). This was in line with our expectations and previous findings, which suggest that donor genetics and other lifestyle factors are the largest source of variability in RNA sequencing samples (Li et al., 2024). This buttresses our approach of performing analysis within matched samples from our donors, where each donor has its own treated and control samples.

**Figure 2.**
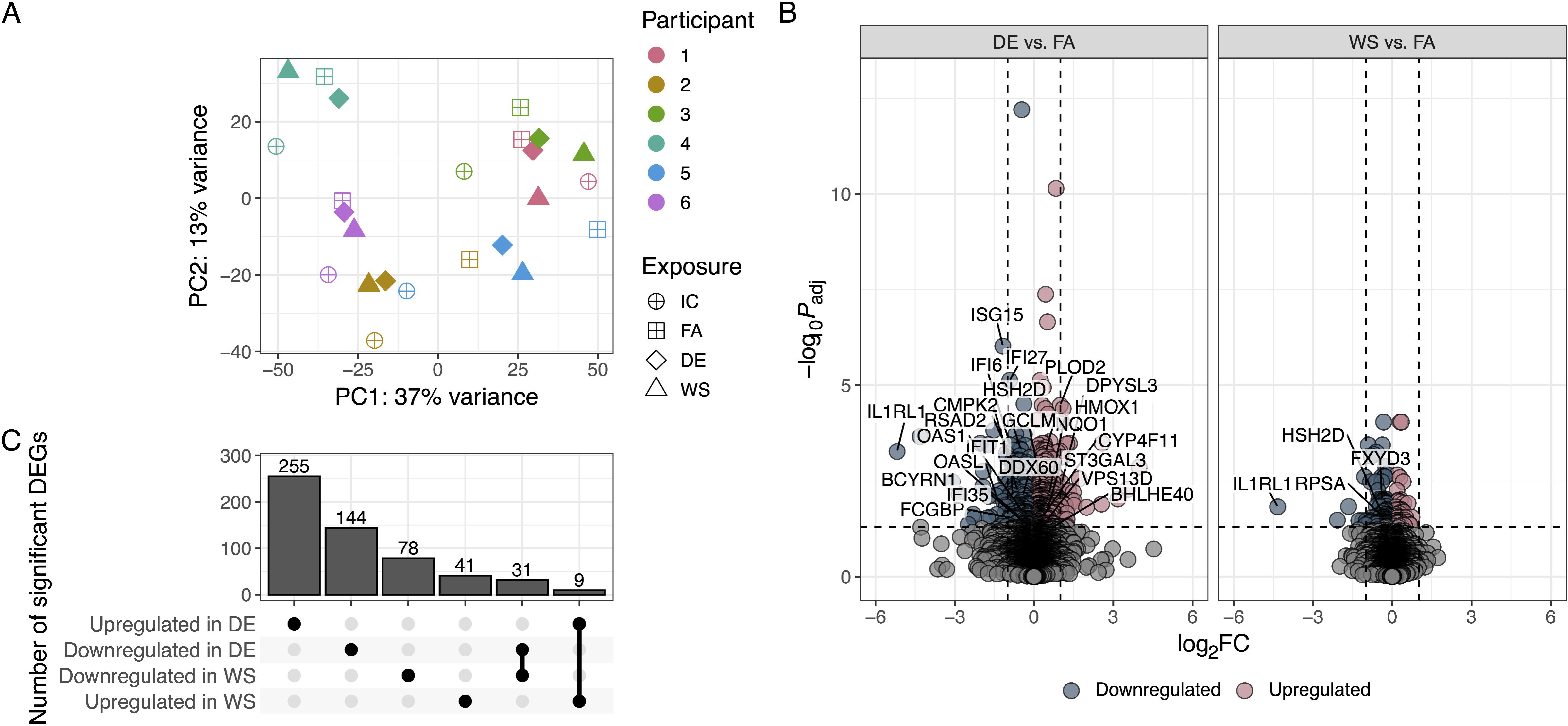
RNA-sequencing analysis of human bronchial epithelial cells (HBECs) exposed to diesel exhaust (DE) or woodsmoke (WS) relative to filtered air (FA). A) Principal component analysis was conducted on the sample count matrix, and PCs 1 and 2 plotted. B) Modelling was used to generate differentially expressed genes (DEGs) for DE and WS in comparison to FA, and the DEGs were plotted as volcano plots with colour indicating those DEGs with adjusted p values ≤0.05 cut-off. Vertical dashed lines indicate fold changes of 1 and −1. C) Upset plot showing the numbers of DEGs that were significantly upregulated or downregulated by DE and WS exposures, relative to FA.

Comparison of FA exposures with IC revealed 3398 (↑2009, ↓1389) differentially expressed genes (DEGs) with the top ten most significant comprising *AC008397.1*, *MRPS15*, *POLB*, *OXSM*, *ABHD11-AS1*, *TFPI2*, *MLX*, *ARL14EP*, *BBX*, and *FXYD3*, at a false discovery rate of <0.05 (Table E3). Notably, the number of dysregulated genes between these two controls potentially reflects mechanical or other stresses experienced during CULTEX exposure, due to airflow over the cells (Buckley et al., 2024). On this basis, we performed all further comparative analyses against cells exposed to FA in the CULTEX, which controls for exposure-associated stress effects.

A total of 159 (↑50, ↓109) DEGs were significantly modulated in WS-exposed cells compared to FA (Table E4). The most significantly modulated DEGs (*CCDC9*, *OXSM*, *TOB2*, *SERTAD1*, *AP003419.1*, *TFPI2*, *WFDC21P*, *TMEM156*, *GPR87*, *ARL14EP*) included a couple with relevance to respiratory disease; *CCDC9*, a gene associated with primary ciliary dyskinesia, and *TFPI2*, which can inhibit serine proteases to reduce extracellular matrix degradation (Lavergne et al., 2021).

DE exposure modulated 439 (↑264, ↓175) DEGs (Table E5) compared to the FA exposure (Figure 2B). The most significant DEGs for DE (*NEK6*, *FTL*, *SLC43A2*, *TKT*, *ISG15*, *SUDS3*, *IFI27*, *TMEM123*, *ATF5*, *PLOD2*), included ferritin light chain (*FTL*) which is involved in iron detoxification (Zarjou et al., 2019), *ISG15* which plays a critical role in host antiviral responses (Perng and Lenschow, 2018), the related interferon alpha-inducible protein 27 (*IFI27*) (Villamayor et al., 2023), and *ATF5*, a transcription factor involved in regulating responses to cellular stresses (Sears and Angelastro, 2017). In addition to DE modulating a greater number of DEGs than WS, the pattern of more transcripts being significantly repressed than induced with WS was reversed.

WS and DE exposures significantly altered 40 common transcripts (↑31, ↓9) (Figure 2C, Table E4, E5). KEGG functional annotations of these genes included immune responses (*IL1RL1*, *SLAMF7*, *CD47)*, mitochondrial function (*COX7B*, *ABHD11*, *MRPS12*) and DNA damage (*DDB1, NUCKS1, AEN*).

### 3.2. Pathway and network effects of exposures

Pathway analysis indicated that WS exposure activated pathways including one associated with Cushing syndrome, while repressing genes related to insulin resistance, adenosine monophosphate-activated protein kinase (AMPK) signaling, and asthma (Figure 3A). Conversely, DE exposure activated pathways including those associated with hypoxia-inducible factor (HIF)-1 signaling, carbon metabolism, and glycolysis/gluconeogenesis, while repressing pathways associated with thyroid hormone synthesis and autoimmune thyroid disease, salivary secretion, and asthma (Figure 3A). When we examined gene networks, we identified small ribosomal subunit and cytochrome complex clusters within the WS DEGs (Figure 3B). In the DE exposure DEGs, we identified changes in a substantially larger network of genes, with clusters relating to respiratory chain complex, the HIF-1 signaling pathway, and a number of genes involved in interferon alpha and beta signaling or interferon-stimulated gene 15 (ISG15)-protein conjugation (Figure 3C).

**Figure 3.**
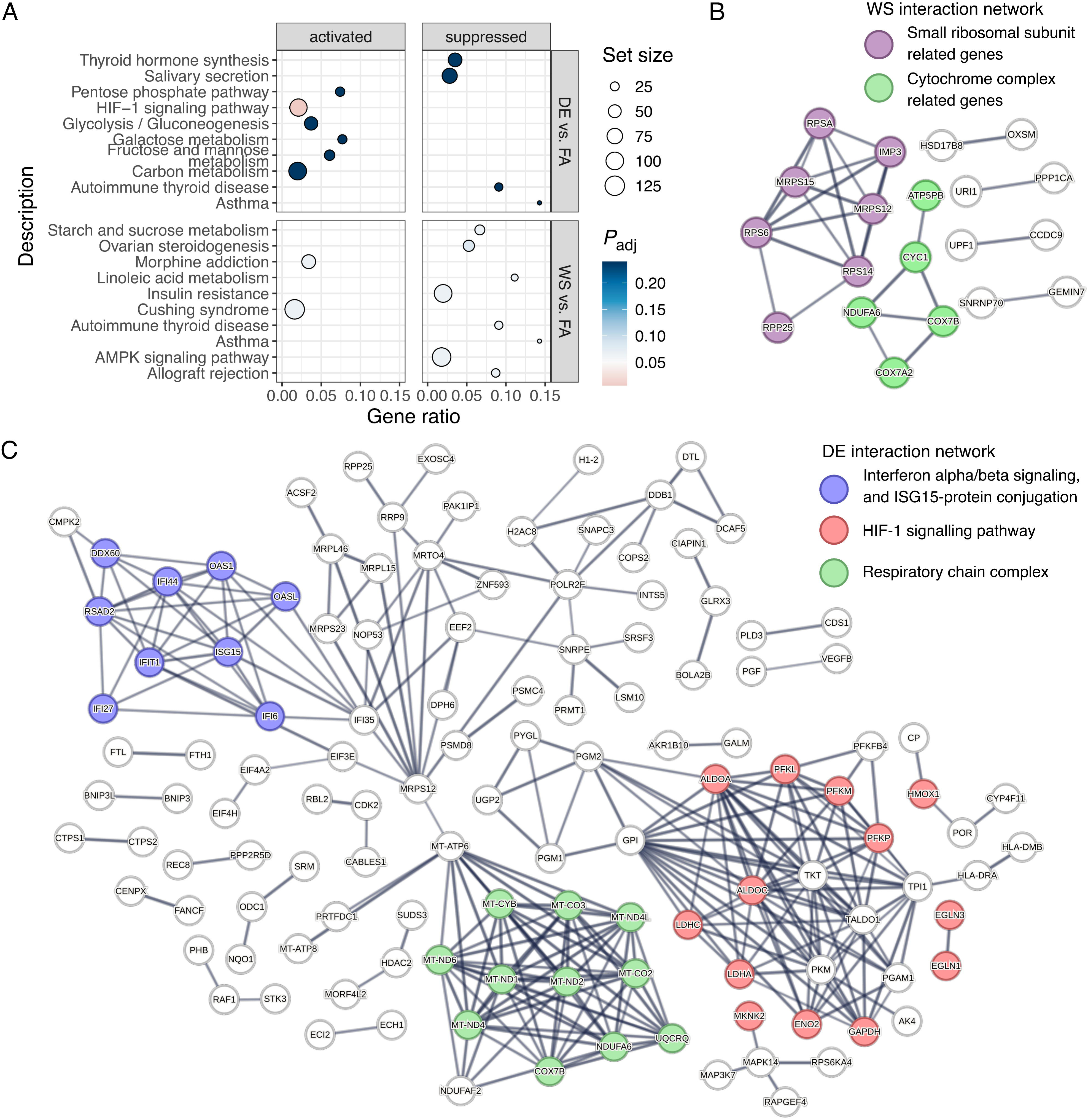
Analysis of enriched pathways. A) Significantly activated or repressed biological processes as identified through GSEA of significantly modulated DEGs. B) Interaction network generated from genes that were differentially expressed upon woodsmoke exposure. As there were no enriched pathways, clusters were annotated based on known functions of genes and coloured accordingly. C) Interaction network generated from genes that were differentially expressed upon diesel exhaust exposure. Coloured nodes represent genes involved in enriched pathways.

Given the network data and to address our a-priori hypotheses, we examined if the DEGs identified for DE and WS exposures were involved in host antiviral responses or oxidative stress responses (see supplemental methods). Analysis of lists of significant DEGs for DE and WS showed that WS repressed four host antiviral response genes (Ioannidis et al., 2012; Mostafavi et al., 2016; Schoggins and Rice, 2011; Troy and Bosco, 2016), while DE modulated 20 (↑6, ↓14) (Table E6). We observed that two known host antiviral response genes, *IL1RL1* and *HSH2D,* were downregulated by both WS and DE (Table E6), while two genes involved in oxidative stress responses, *DKK1* and *NF2,* were repressed and induced, respectively, in both WS and DE-exposed cells (Tables E4 and E5). DE also repressed several genes negatively regulated by the oxidative stress-responsive transcription factor Nrf2 (*NFE2L2*), such as *RSAD2, IFIT1*, and *ISG15* (Robertson et al., 2020). To explore these responses, we considered 12 specific genes previously identified within the Nrf2 and aryl hydrocarbon receptor (AHR) pathways (Huff et al., 2020). Of these, *GCLM, NQO1,* and *HMOX1* were upregulated only by DE (Table E5).

### 3.3. Effects of diesel exhaust and wood smoke exposures on cytokines

As a comprehensive analysis of the effects of exposure to DE and WS, we also evaluated the changes in a protein immune-mediators panel. Consistent with our RNAseq analysis, samples clustered by donor (Figure E3). We also examined overall differences in mediator concentrations across all conditions (Figure 4A). Immune mediator profiles were mostly consistent between IC and FA conditions, with 5 of 59 mediators differing significantly. When examining FA and DE, significant differences were observed for 12 mediators compared to only one mediator (mutually exclusive) for FA vs WS (Figure 4A). We also observed DE had an overall suppressive effect on immune mediator levels.

**Figure 4.**
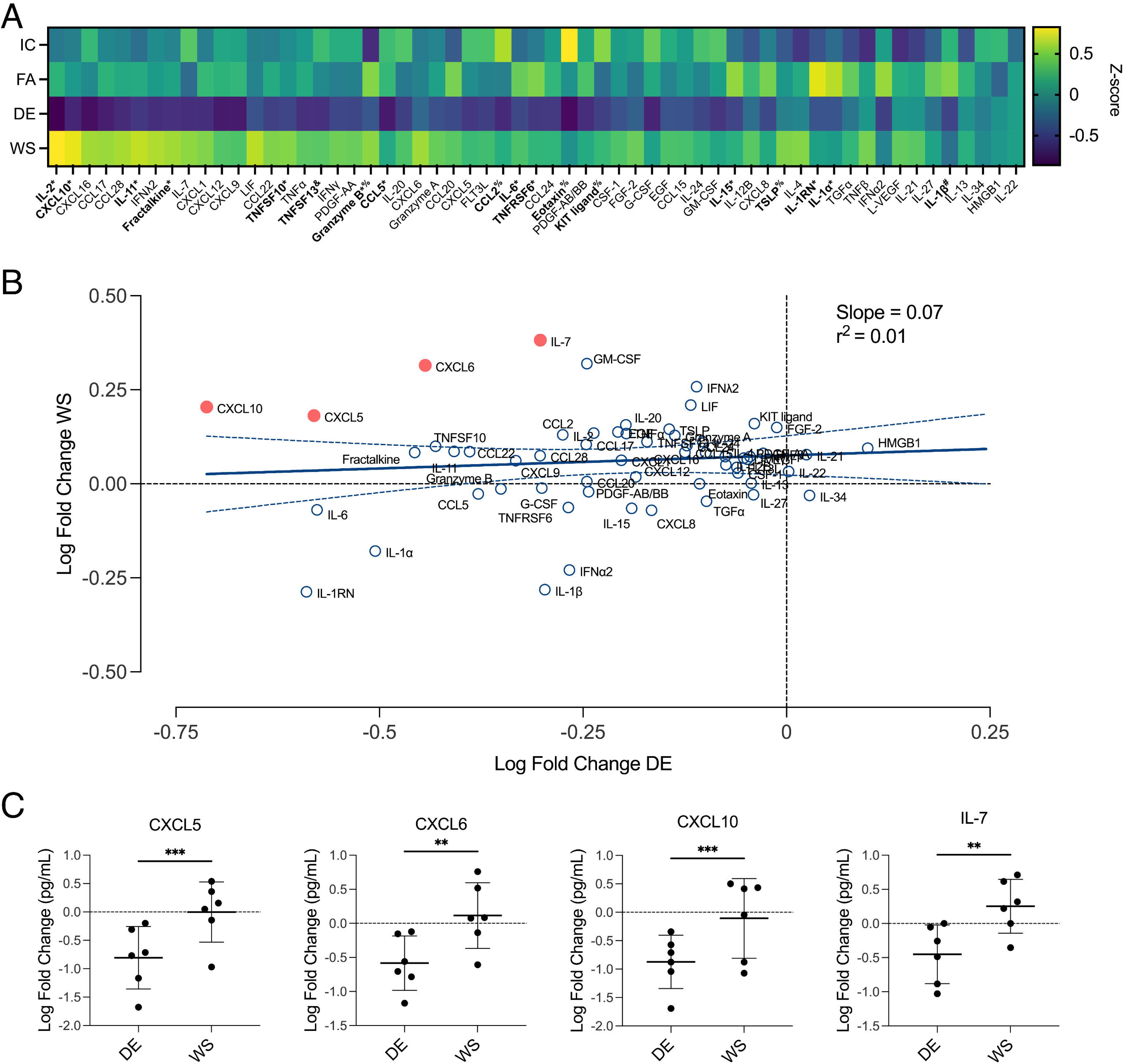
A) Heatmap of cytokine responses following IC, FA, DE and WS exposures. Z-scores shown ranked in descending order of the difference between WS and DE. Concentration (pg/mL) data were log-transformed and analyzed by two-way ANOVA with Tukey Post-Hoc Test (n = 6). Significant differences (p < 0.05): *DE vs FA, ^#^WS vs FA, ^&^DE vs WS. ^%^IC vs FA. B) Log fold change of immune mediators (pg/mL) for WS and DE versus FA was plotted. Linear regression was used to estimate correlation. FA-normalized data was analyzed using a two-way ANOVA with Šidák Post-Hoc Test and mediators significantly different between WS and DE are highlighted in red. C-G) Log fold change of immune mediators (pg/mL) for WS and DE versus FA was plotted. FA-normalized data was analyzed using a two-way ANOVA with Šidák Post-Hoc Test. Significant differences (p < 0.01**, p < 0.001**).

To better understand the differences observed between DE and WS, both conditions were normalized to FA and then compared (Figure 4B). Indeed, immune mediator levels were not correlated between WS and DE-exposed cells (r^2^=0.01). Moreover, neutrophil chemotaxis mediators CXCL5 and CXCL6 and host antiviral response-associated proteins CXCL10 and IL-7 significantly differed between WS and DE, with a suppressive effect observed in DE relative to WS-exposed cells (Figure 4B and C).

## 4. Discussion

We demonstrate that two major sources of climate change-related air pollution modulate different genes sets and pathways in human bronchial epithelial cells. Both DE and WS exposures dysregulated 40 common genes involved in immune responses, mitochondrial function and DNA damage. Of note, the overlap of dysregulated gene expression between the two air pollutants was modest, suggesting distinct responses of the airway epithelium to these pollutants. At similar controlled PM_2.5_ concentrations, DE induced more DEGs than WS and more DEGs were upregulated by DE than repressed, while WS showed the opposite pattern. Our findings differ from previous studies using A549 or BEAS-2B cell lines, which demonstrated significant increases in CXCL8 with both exposures (Wang et al., 2021; Zheng et al., 2015). These discrepancies may arise from differences in cellular composition and cell source; our model of primary HBECs in ALI better reflects the *in vivo* airway epithelium. Thus, our results speak to the importance of performing controlled exposure experiments on primary cells, more representative of human airways, to assess the toxicity of exposures.

Our findings show that DE exposure repressed more host antiviral-response DEGs and proteins than WS, suggesting that TRAP exposure may increase infection risk to a greater degree. Data increasingly supports the link between air pollution and increased prevalence and severity of viral infections (Domingo and Rovira, 2020; Monoson et al., 2023; Rebuli et al., 2021; Troy and Bosco, 2016). However, WS significantly altered immunity-related transcripts, belying its perception as a simple aesthetic enhancement, as reflected in the popularity of fireplaces and campfires. Few studies (and no controlled human exposure studies (Long et al., 2024)) compare WS to pollution types like TRAP, a gap our paper addresses.

The cause of the differential transcriptomic responses elicited by DS and WS exposures in our study is unclear. Analysis of our DE and WS exposures at matched PM_2.5_ mass concentrations revealed differences in gasses present (Table E2). Specifically, the levels of NO, CO, CO_2_, and O_3_ were all elevated in DE exposures compared to WS exposures. Previous studies have examined the effect of specific gaseous components of air pollution such as NO and O_3_ and found increases in genes related to antioxidant defense, oxidative stress, and inflammation in exposed HBECs (Bayarri et al., 2021; Bromberg, 2016). One study, in particular, looked at the acute effects of a 2-hour exposure to O_3_ on a panel of genes related to cellular toxicity and oxidative stress, and reported 60 differentially expressed genes (Mirowsky et al., 2016). In our study we observed significant differences in four of these genes (*FTH1*, *HMOX1*, *SLC2A1*, and *BNIP3*) in DE and none of these genes in WS exposed cells, indicating that the differential levels of gas in our exposures may partially explain the differences we observed in transcriptome profiles. However, it is important to note that the chemical composition of the PM also likely differed and influenced HBEC responses in our system. Further characterization and chemical analysis of the exposure PM may help explain differences in HBEC responses.

Our study has laid out a framework for assessing the transcriptomic and cytokine changes due to air pollution exposure. Important features of the system include particle-level controlled exposure to air pollutants at equivalent mass concentration, amenability to human primary cell culture, and ease of collection of cells and supernatants for molecular profiling. Thus, our framework can be readily applied to future controlled exposure experiments involving different air pollutants and different cell or tissue types, and can accommodate studies testing measures to mitigate the molecular effects of exposures.

Nevertheless, our work has some limitations. The 300 µg/m³ PM_2.5_ concentration we used does not reflect the full range of real-world levels, but parallels typical controlled human exposure study concentrations and also approximates levels intermittently present in polluted Asian cities and hotspots across North America, such as are experienced during commutes (Carlsten et al., 2016; Luglio et al., 2024; Ryu et al., 2022; Wooding et al., 2019). Additionally, while 24 hours may not be optimal for detecting initial transcriptional effects after exposure, this timepoint was chosen based on the relevant literature and to optimize our ability to detect cytokine responses in parallel. A majority of similar studies of cell exposures to DE or WS conducted to date have also utilized this 24-hour exposure time (Holder et al., 2008; Knebel et al., 2002; Mullins et al., 2016; Zarcone et al., 2016). Finally, WS alone is a limited proxy for wildfires, though an excellent model of residential wood burning for heating or cooking which also contributes significantly to outdoor air pollution (Hong et al., 2017; Languille et al., 2020; Ward et al., 2017). We chose to use lodgepole pine as it is a standard wood type used by the US EPA in similar experiments, and Canada’s largest national firewood distributors, and given its prevalence in Western Canadian forests (Bull and Bennett, 2022; Gagné et al., 2017; Ministry of Forests, 2024). However, our system could be used to burn any wood type or other relevant combustible matter. However, wildfires have substantial compositional variation and complex interactions with factors like weather, temperature, humidity, and sunlight, which can also affect atmospheric aerosol aging and secondary organic aerosol production, making it essentially impossible to accurately model them in a lab setting. We, therefore, caution against generalizing the relative toxicity of WS and DE until further *in vitro* and human exposure studies are completed.

Although it is beyond the scope of this manuscript, further investigation is needed to understand whether underlying disease and/or genetic susceptibility modifies cellular and tissue responses to these air pollutants and how gene-environment interactions are different based on the source of air pollutants. Such research would allow risk stratification for those with increased susceptibility to air pollutants (Carlsten et al., 2014; Laumbach et al., 2021). Furthermore, *in vitro* systems such as ours, when combined with model systems perturbing the genome (e.g., CRISPR-based genetic screening Perturb-seq (Dixit et al., 2016)), would allow direct investigation of gene variant-environment interaction, which is not possible with *in vivo* human studies.

Nonetheless, our data informs the debate regarding whether current regulatory focus on concentration should additionally consider composition, along with the related assumption that similar concentrations of air pollution have indistinct effects. Our findings emphasize the need to leverage emerging techniques and systems biology to better understand complex pollution-induced molecular changes to airway health.

## CRediT authorship contribution statement

Conceptualization: RDH, CFR, CC; Data curation: RDH, CFR, TL, MHR, ELL; Formal Analysis: RDH, CFR, TL, KIH, MHR, ELL; Funding acquisition: RDH, CFR, CC; Investigation: RDH, NE; Methodology: RDH, CFR, TL, MHR, CC, ELL; Project administration: RDH, CFR, MHR, CC, ELL; Resources: RDH, TL, NE, MHR, CC, ELL; Software: RDH, CFR, TL, KIH, MHR, ELL; Supervision: CFR, CC, ELL; Validation: RDH, CFR, TL, MHR, ELL; Visualization: RDH, CFR, TL, MHR, ELL; Writing – original draft: RDH, CFR, TL, MHR, CC, ELL; Writing – review & editing: RDH, CFR, TL, KIH, NE, MHR, CC, ELL.

## Acknowledgement

We want to thank our COPA and AGnES study participants, without whom this research would not have been possible. We are grateful to Carley Schwartz who, with significant guidance from Drs. Ian Gilmour and YongHo Kim, developed the woodsmoke exposure system. We are also grateful to Yu Xi, Agnes Yuen, PJ, Tina Afshar, Denise Wooding, Hang Li, Juma Orach, Kevin Lau, Illiassou Hamidou and everyone else involved in completing the AGnES and COPA studies, and this work. We would also like to thank our close collaborators Drs. Emily Brigham, Kelly McNagny, Neeloffer Mookherjee, Neil Alexis and Meghan Rebuli for their support of our quest to find related funding for a human exposure study. The authors also wish to acknowledge Canada’s Michael Smith Genome Sciences Centre, Vancouver, Canada for completing library construction and sequencing. A full list of funders of infrastructure and research supporting the Michael Smith Genome Sciences Centre services accessed is available at www.bcgsc.ca/about/funding_support. This research was conducted on Musqueam, Squamish, and Tsleil-Waututh traditional territory.

## Data and code availability

All data are available on the European Genome-phenome Archive (https://ega-archive.org/). All scripts used for analyses can be found on GitHub (https://github.com/EmiliaLimLab/DE_WS_RNAseq).

## Use of artificial intelligence

During the preparation of this work the authors used Microsoft Copilot to provide suggestions for streamlining the text. The authors edited all text manually and take full responsibility for the content of the publication.

## Supplementary Figure Legends

**Figure E1.**
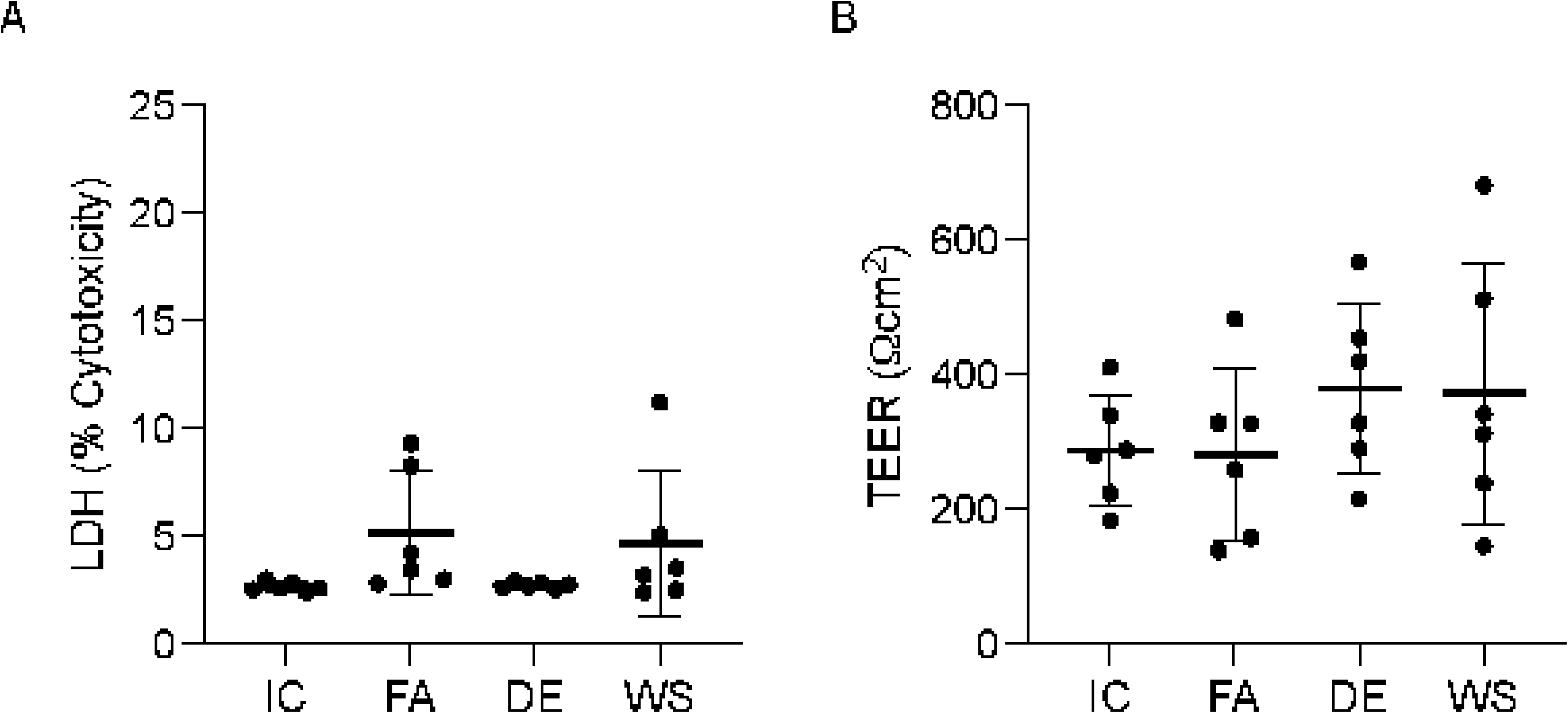
Cytotoxicity and trans-epithelial electrical resistance (TEER) for each condition. LDH assays (A) were performed using basolateral supernatants collected at 24 hours. The percent cytotoxicity (percent of lysis buffer control) is plotted as mean ± SD. (B) TEER was measured 24 hours post-exposure in Hank’s balanced salt solution and is plotted as mean ± SD.

**Figure E2.**
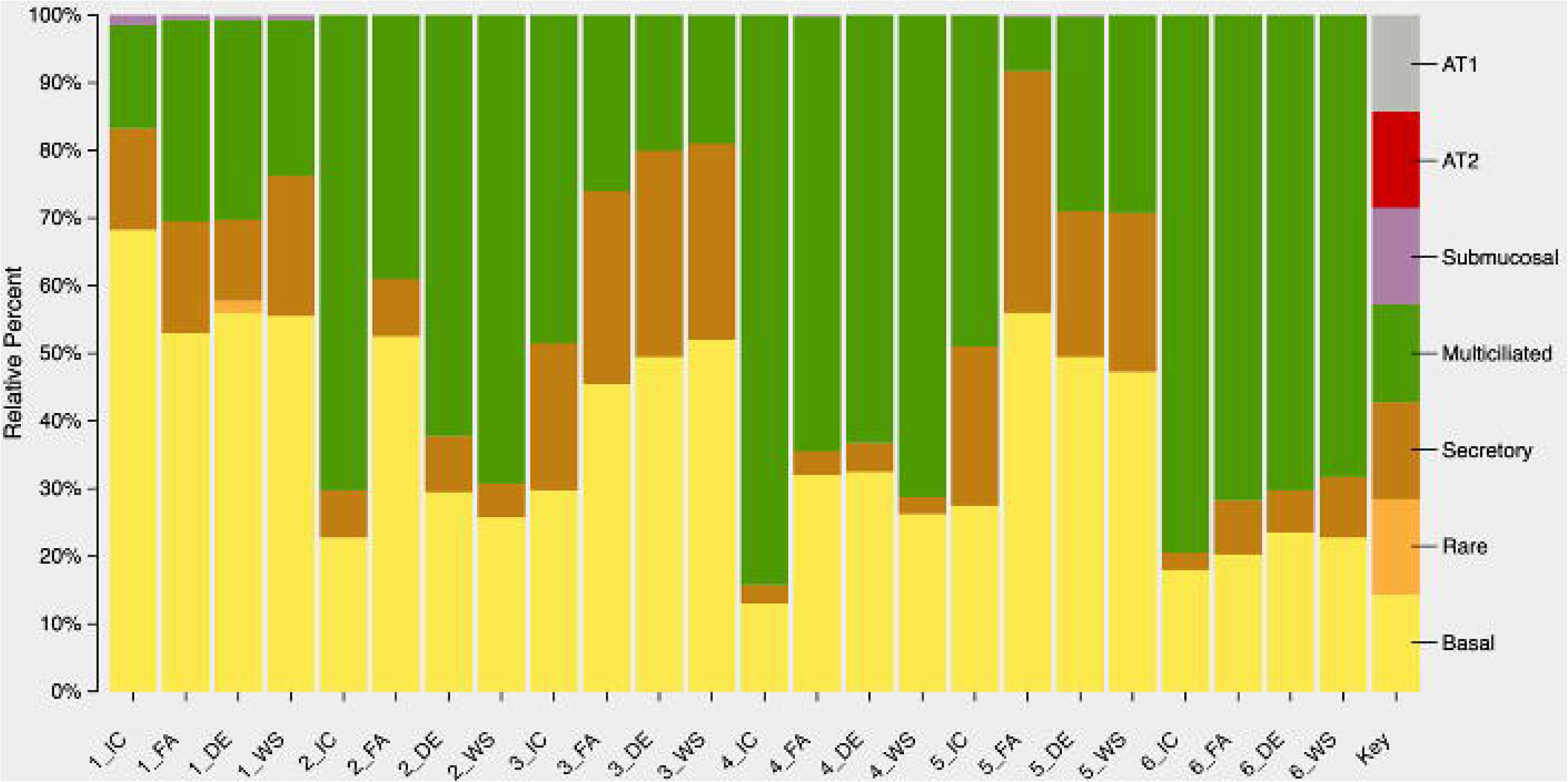
CIBERSORTx imputation of epithelial cell types. CIBERSORTx (https://cibersortx.stanford.edu), calibrated on Human Lung Cell Atlas data, was used to impute the proportions of basal and other epithelial cell types from the count matrix values (CPM values) of each sample. Relative percent proportions for basal, rare, secretary, multiciliated lineage, submucosal secretary, AT1 and AT2 cells were plotted for each sample.

**Figure E3.**
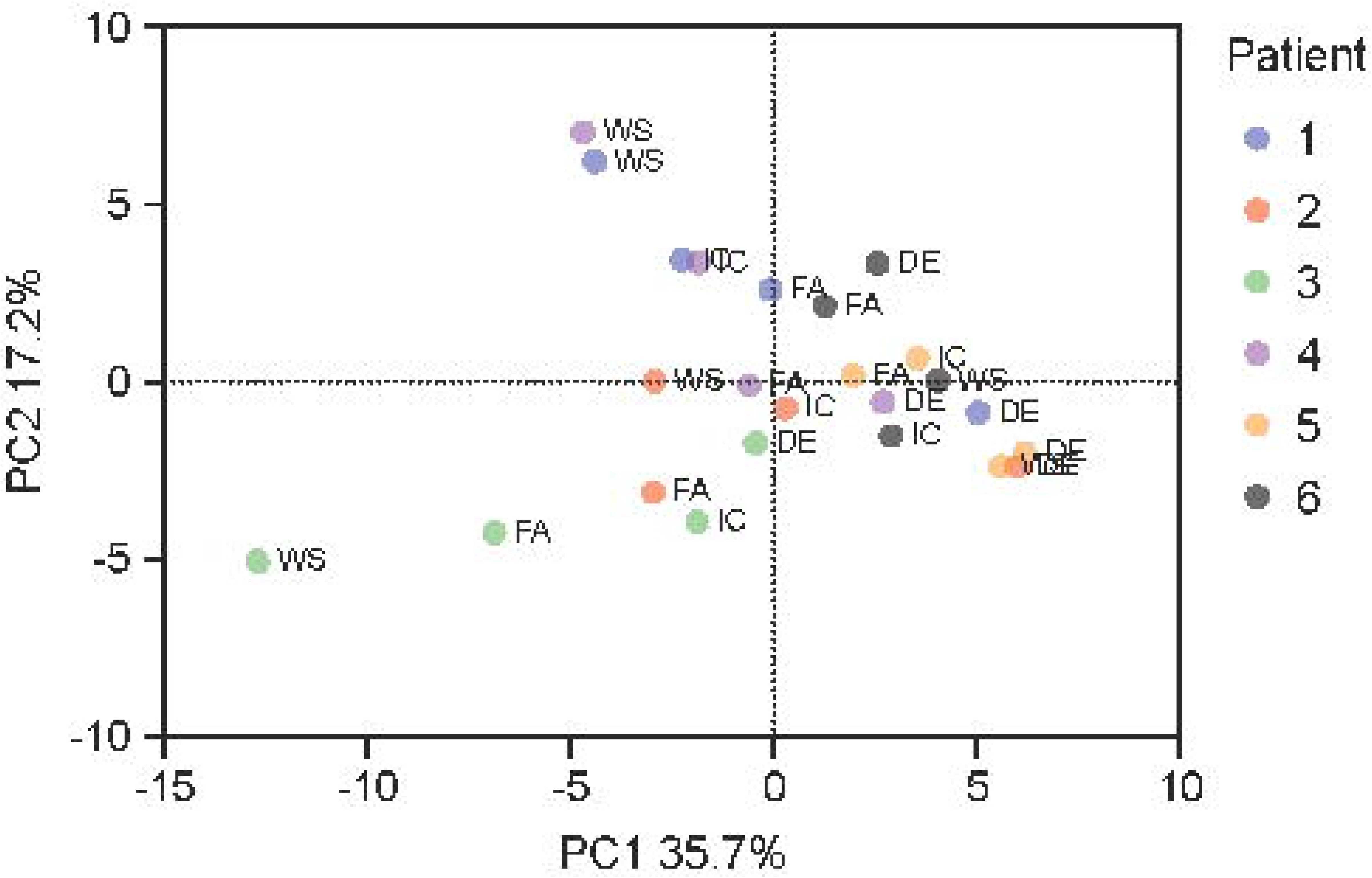
PCA of cytokine responses following IC, FA, DE and WS exposures. Principal component analysis was conducted on the data received from the immune mediator protein analysis, and principal components 1 and 2 plotted as a scatterplot.

## Supplementary Table Legends

**Table E1.** Demographics of human bronchial epithelial cell (HBEC) donors. Bronchial epithelial cells were collected during research bronchoscopies conducted for the AGnES and COPA studies (see methods). Donors with a modest smoking history had not smoked for at least 6 months prior to HBEC collection. FEV_1_, forced expiratory volume in 1 second; FVC, forced vital capacity.

**Table E2.** Exposure characteristics of the human bronchial epithelial (HBEC) cells. Air liquid interface cultured HBECs collected from six study participants (Table E1) were exposed for 2-hours to either diesel exhaust or wood smoke each diluted and standardized to a nominal PM_2.5_ concentration of 300 µg/m^3^. Post-dilution concentrations of gasses and other aerosol characteristics are also reported. Standard deviation (SD) shows variability of measurements over the 2-hour exposure. CO, carbon monoxide; CO_2_, carbon dioxide; NO, nitric oxide; NO_2_, nitrogen dioxide; O_3_, ozone; PM_2.5_, particulate matter with a diameter of less than 2.5 micrometers; ppb, parts per billion; ppm, parts per million; TVOC, total volatile organic compounds.

**Table E3**. Significantly modulated DEGs following filtered air (FA) exposure in comparison to incubator control (IC). Also available on our GitHub page (https://github.com/EmiliaLimLab/DE_WS_RNAseq).

**Table E4**. Significantly modulated DEGs following woodsmoke (WS) exposure in comparison to filtered air (FA). Also available on our GitHub page (https://github.com/EmiliaLimLab/DE_WS_RNAseq).

**Table E5**. Significantly modulated DEGs following diesel exhaust (DE) exposure in comparison to filtered air (FA). Also available on our GitHub page (https://github.com/EmiliaLimLab/DE_WS_RNAseq).

**Table E6.** Host antiviral response-related DEGs repressed following DE and/or WS exposure. A list of relevant genes was extracted from the literature (Ioannidis et al., 2012; Mostafavi et al., 2016; Schoggins and Rice, 2011; Troy and Bosco, 2016) and imported into R. The gene list was compared with our lists of DE and WS exposure-modulated DEGs using the VennDetail R package. DEGs that were found on both our diesel exhaust (DE) and woodsmoke (WS) significantly modulated list that matched with the literature gene list are plotted in the first section, followed by those only matching with DE in the second section, and finally those only matching with WS at the bottom.

