## Supplemental Methods for "Woodsmoke and Diesel Exhaust: Distinct Transcriptomic Profiles in the Human Airway Epithelium"

Full length article

Running title: Transcriptomics of woodsmoke or diesel exhaust

Keywords: Gene expression, Air pollution, RNA sequencing, Oxidative stress pathways, Host antiviral responses

Corresponding author at:

Air Pollution Exposure Lab, University of British Columbia,

2775 Laurel St. 7^th^ Floor, the Lung Center, Vancouver General Hospital – Gordon and Leslie Diamond Health Care Centre, Vancouver, BC, V5Z 1M9, Canada

**S1. Supplemental Methods:**

*S1.1. Detailed methods for RNA sequencing*

Note that this RNA-sequencing protocol was completed and authored by Michael Smith Genome Science Centre (Vancouver, Canada) personnel

Qualities of total RNA samples were determined using an Agilent Bioanalyzer RNA Nanochip. Polyadenylated (PolyA+) RNA was purified using the NEBNext Poly(A) mRNA Magnetic Isolation Module (E7490L, New England Biolabs) from 35-660 ng total RNA. Messenger RNA selection was performed using NEBNext Oligod(T)25 beads (NEB) incubated at 65 °C for 5 minutes followed by snap-chilling at 4 °C to denature RNA and facilitate binding of poly(A) mRNA to the beads. mRNA was eluted from the beads in Tris Buffer incubated at 80 °C for 2 minutes then held at 25 °C for 2 minutes. RNA binding buffer was added to allow the mRNA to re-bind to the beads, mixed 10 times and incubated at room temperature for 5 minutes. The sample plate was placed on the magnet and the supernatant discarded. The mRNA bound beads were washed twice then cleared again on magnet. The supernatant was again discarded and mRNA eluted from the beads in 20 µL Tris buffer incubated at 80 °C for 2 minutes. mRNA was transferred to a new 96-well plate.

First-strand cDNA was synthesized from heat-denatured purified mRNA using a Maxima H Minus First Strand cDNA Synthesis kit (Thermo-Fisher, USA) and random hexamer primers at a concentration of 200 ng/µL along with a final concentration of 40 ng/µL Actinomycin D, followed by PCR Clean DX (Aline Biosciences) bead purification on a Microlab NIMBUS robot (Hamilton Robotics, USA). The second strand cDNA was synthesized following the NEBNext Ultra Directional Second Strand cDNA Synthesis protocol (New England Biolabs) that incorporates dUTP in the dNTP mix, allowing the second strand to be digested using USERTM enzyme (NEB) in the post-adapter ligation reaction and thus achieving strand specificity.

cDNA was fragmented to by Covaris LE220 sonication to achieve 250-300 bp average fragment lengths. The paired-end sequencing library was prepared following the BC Cancer Genome Sciences Centre strand-specific, plate-based library construction protocol on a Microlab NIMBUS robot (Hamilton Robotics, USA). Briefly, the sheared cDNA was subject to end-repair and phosphorylation in a single reaction using an enzyme premix (New England Biolabs) containing T4 DNA polymerase, Klenow DNA Polymerase and T4 polynucleotide kinase, incubated at 20 °C for 30 minutes. Repaired cDNA was purified in 96-well format using PCR Clean DX beads (Aline Biosciences, USA), and 3’ A-tailed (adenylation) using Klenow fragment (3’ to 5’ exo minus) and incubation at 37 °C for 30 minutes prior to enzyme heat inactivation. Illumina TruSeq adapters were ligated at 20 °C for 15 minutes. The adapter-ligated products were purified using PCR Clean DX beads, then digested with USERTM enzyme (1U/µL, NEB) at 37^o^C for 15 minutes followed immediately by 10 cycles of indexed PCR using NEBNext Ultra II Q5 Master Mix (New England Biolabs) and Illumina’s primer set. PCR parameters: 98 ^o^C for 1 minute followed by 10 cycles of 98 ^o^C for 15 seconds, 65 ^o^C for 30 seconds and 72 ^o^C for 30 seconds, and then 72 ^o^C for 5 minutes. The PCR products were purified and size-selected using a 1:1 PCR Clean DX beads-to-sample ratio (twice), and the eluted DNA quality was assessed with Caliper LabChip GX for DNA samples using the High Sensitivity Assay (PerkinElmer, Inc. USA) and quantified using a Quant-iT dsDNA High Sensitivity Assay Kit on a Qubit fluorometer (Invitrogen) prior to library pooling and size-corrected final molar concentration calculation for Illumina sequencing with paired-end 150 base reads.NA samples were determined using an Agilent Bioanalyzer RNA Nanochip. Polyadenylated (PolyA+) RNA was purified using the NEBNext Poly(A) mRNA Magnetic Isolation Module (E7490L, New England Biolabs) from 35-660 ng total RNA. Messenger RNA selection was performed using NEBNext Oligod(T)25 beads (NEB) incubated at 65 °C for 5 minutes followed by snap-chilling at 4 °C to denature RNA and facilitate binding of poly(A) mRNA to the beads. mRNA was eluted from the beads in Tris Buffer incubated at 80 °C for 2 minutes then held at 25 °C for 2 minutes. RNA binding buffer was added to allow the mRNA to re-bind to the beads, mixed 10 times and incubated at room temperature for 5 minutes. The sample plate was placed on the magnet and the supernatant discarded. The mRNA bound beads were washed twice then cleared again on magnet. The supernatant was again discarded and mRNA eluted from the beads in 20 µL Tris buffer incubated at 80 °C for 2 minutes. mRNA was transferred to a new 96-well plate.

First-strand cDNA was synthesized from heat-denatured purified mRNA using a Maxima H Minus First Strand cDNA Synthesis kit (Thermo-Fisher, USA) and random hexamer primers at a concentration of 200 ng/µL along with a final concentration of 40 ng/µL Actinomycin D, followed by PCR Clean DX (Aline Biosciences) bead purification on a Microlab NIMBUS robot (Hamilton Robotics, USA). The second strand cDNA was synthesized following the NEBNext Ultra Directional Second Strand cDNA Synthesis protocol (New England Biolabs) that incorporates dUTP in the dNTP mix, allowing the second strand to be digested using USERTM enzyme (NEB) in the post-adapter ligation reaction and thus achieving strand specificity.

cDNA was fragmented to by Covaris LE220 sonication to achieve 250-300 bp average fragment lengths. The paired-end sequencing library was prepared following the BC Cancer Genome Sciences Centre strand-specific, plate-based library construction protocol on a Microlab NIMBUS robot (Hamilton Robotics, USA). Briefly, the sheared cDNA was subject to end-repair and phosphorylation in a single reaction using an enzyme premix (New England Biolabs) containing T4 DNA polymerase, Klenow DNA Polymerase and T4 polynucleotide kinase, incubated at 20 °C for 30 minutes. Repaired cDNA was purified in 96-well format using PCR Clean DX beads (Aline Biosciences, USA), and 3’ A-tailed (adenylation) using Klenow fragment (3’ to 5’ exo minus) and incubation at 37 °C for 30 minutes prior to enzyme heat inactivation. Illumina TruSeq adapters were ligated at 20 °C for 15 minutes. The adapter-ligated products were purified using PCR Clean DX beads, then digested with USERTM enzyme (1U/µL, NEB) at 37 °C for 15 minutes followed immediately by 10 cycles of indexed PCR using NEBNext Ultra II Q5 Master Mix (New England Biolabs) and Illumina’s primer set. PCR parameters: 98 °C for 1 minute followed by 10 cycles of 98 °C 15 seconds, 65 °C 30 seconds and 72 °C 30 seconds, and then 72 °C 5 minutes. The PCR products were purified and size-selected using a 1:1 PCR Clean DX beads-to-sample ratio (twice), and the eluted DNA quality was assessed with Caliper LabChip GX for DNA samples using the High Sensitivity Assay (PerkinElmer, Inc. USA) and quantified using a Quant-iT dsDNA High Sensitivity Assay Kit on a Qubit fluorometer (Invitrogen) prior to library pooling and size-corrected final molar concentration calculation for Illumina sequencing with paired-end 150 base reads. Sequencing was performed with an Illumina NovaSeq 6000 sequencer targeting 50M read-pairs per library.

*S1.2. Oxidative stress pathways and host antiviral response analysis*

Based on our a-priori hypotheses supported by initial analyses of DEGs and KEGG pathways, we identified oxidative stress transcription factor pathways and associations with host antiviral responses as areas to evaluate further. We sought to compare significant DEGs from our DE and WS exposures, relative to FA, to lists of genes regulated by the oxidative stress response transcription factors nuclear factor erythroid 2-related factor 2 (*NFE2L2*/Nrf2) and the aryl hydrocarbon receptor (AHR). Lists of genes regulated by Nrf2 and AHR were extracted from the TF2DNA database hosted by the Fiser lab at <http://fiserlab.org/tf2dna_db/>, with the Kulakovskiy2013 data extracted (Kulakovskiy et al., 2013). To provide a comprehensive list of targets of these transcription factors, the TFlink database at <https://tflink.net/> was also consulted and the *Homo sapiens* lists for NFE2L2 (<https://tflink.net/protein/q16236/>) and AHR (<https://tflink.net/protein/p35869/>) extracted. The TF2DNA and TFlink data was imported into R (v4.4.2), and the gene lists merged using the VennDetail R package (v1.20.0).

We also consulted literature on host antiviral responses (Ioannidis et al., 2012; Mostafavi et al., 2016; Schoggins and Rice, 2011; Troy and Bosco, 2016) and a list of significant DEGs from primary airway epithelial cells infected with influenza or respiratory syncytial virus (Ioannidis et al., 2012). Lists of genes, including those from Ioannidis et al. were extracted and imported into R for further analysis. The lists of genes for Nrf2, AHR and host antiviral responses were compared with our DEG lists using the VennDetail R package. The results of these analyses of Nrf2/AHR and host antiviral responses are reported within the main text of our paper and in Table E7, respectively.
